# Anti-GD2 antibody therapy alters the neuroblastoma tumor microenvironment and extends survival in *TH-MYCN* mice

**DOI:** 10.1101/2021.06.09.447746

**Authors:** KO McNerney, S Karageorgos, G Ferry, A Wolpaw, C Burudpakdee, P Khurana, R Vemu, A Vu, MD Hogarty, H Bassiri

**Author notes:** These authors contributed equally to this work (KO McNerney, S Karageorgos). These authors contributed equally to this work (MD Hogarty, H Bassiri). SK current affiliation is First Department of Pediatrics, National and Kapodistrian University of Athens, “Aghia Sophia” Children’s Hospital, Athens, Greece. GF current affiliation is University College London, Great Ormond Street, Institute of Child Health. Correspondence: Michael D. Hogarty, Hamid Bassiri, Addresses: Michael Hogarty, CTRB, Room 3020, 3501 Civic Center Blvd, Philadelphia, PA 19104, Hamid Bassiri, CTRB, Room 3100, 3501 Civic Center Blvd, Philadelphia, PA 19104.

## Abstract

**Background:** Neuroblastoma is a commonly lethal solid tumor of childhood and intensive chemoradiotherapy treatment cures ~50% of children with high-risk disease. The addition of immunotherapy using dinutuximab, a monoclonal antibody directed against the GD2 disialoganglioside expressed on neuroblasts, improves survival when incorporated into front-line therapy and shows robust activity in regressing relapsed disease when combined with chemotherapy. Still, many children succumb to neuroblastoma despite receiving dinutuximab-based immunotherapy, and efforts to counteract the immune suppressive signals responsible are warranted. Animal models of human cancers provide useful platforms to study immunotherapies. *TH-MYCN* transgenic mice are immunocompetent and develop neuroblastomas at autochthonous sites due to enforced *MYCN* expression in developing neural crest tissues. However, GD2-directed immunotherapy in this model has been underutilized due to the prevailing notion that *TH-MYCN* neuroblasts express insufficient GD2 to be targeted.

**Methods:** *TH-MYCN* mice were treated with 14G2a (anti-GD2 antibody), isotype antibody, or phosphate buffered saline from day 14 of life until day 100 or signs of morbidity. Survival was recorded, and tumors were isolated in terminal surgeries for analysis of GD2 expression and immune cell frequencies. Tumors from untreated mice were explanted for generation into cell lines, and GD2 expression was recorded with serial passage in tissue culture. Immunocytology and immunoblotting were performed to evaluate for adrenergic and mesenchymal markers of neuroblasts. Survival curves compared using Kaplan-Meier method with a log-rank test for significance. Unpaired two-tailed Student’s *t*-tests used for comparison of groups in flow cytometry analysis.

**Results:** 14G2a markedly extends survival in such *TH-MYCN* mice. Additionally, neuroblasts in 14G2a-treated mice have reduced GD2 expression and fewer macrophage and myeloid-derived suppressor cells in their tumor microenvironments. Neuroblasts in *TH-MYCN*-driven tumors express GD2 at levels comparable to human neuroblastomas but rapidly lose GD2 expression when explanted ex vivo to establish tumor cell lines. The loss of GD2 expression ex vivo is associated with a transition from an adrenergic to mesenchymal state that is maintained when reimplanted in vivo.

**Conclusions:** Our findings support the utility of the *TH-MYCN* model to inform GD2-directed immunotherapy approaches for neuroblastoma as well as opportunities to investigate drivers of adrenergic to mesenchymal fate decisions.

## INTRODUCTION

A variety of immunotherapies are altering the treatment landscape for many cancers, particularly those of hematologic origin. Unfortunately, solid tumor immunotherapies are frequently hindered by the capacity of the tumor microenvironment (TME) to suppress and exclude immune effector cells. Neuroblastoma is a common solid tumor of childhood that carries a 5-year mortality rate of ~50% in its high-risk form despite the use of intensive treatment with surgery, high-dose chemotherapy with stem cell rescue, radiotherapy and biotherapy with retinoids Like most pediatric cancers, neuroblastomas manifest a low mutation burden and a paucity of neo-antigens for the immune system to target. [1–3] Further, its TME is enriched with immune and stromal cells that include tumor associated macrophages (TAMs), myeloid-derived suppressor cells (MDSCs), T-regulatory cells, and fibroblasts that collectively release immunosuppressive cytokines and attenuate host immune responses. [4–6]

Despite these barriers, dinutuximab, a chimeric antibody that targets the GD2 disialoganglioside highly expressed on nearly all neuroblastomas, has shown efficacy when combined with immunostimulatory cytokines in front-line maintenance therapy for patients with high-risk neuroblastoma, leading to its FDA approval, and when used in combination with chemotherapy to treat relapsed or refractory neuroblastoma. [7–9] Although these clinical successes substantiate the promise of immunotherapy for neuroblastoma, overall survival remains low, and many children die from tumor progression despite receiving dinutuximab. A more thorough understanding of the immunosuppressive TME should provide opportunities to improve this therapy. Such mechanistic studies can be achieved in faithful mouse models of cancer, enabling the discovery and preclinical prioritization of novel therapeutic approaches, including combination therapies with GD2-directed antibodies. Despite differences between the murine and human immune systems, immunocompetent transgenic mouse models have been credentialed for the study of immunotherapies and a neuroblastoma model that uses a cancer-relevant oncogenic driver to initiate tumors in immunocompetent mice is paramount to further leverage the activity of dinutuximab. [10–12]

The *Tyrosine-Hydroxylase-MYCN* (*TH-MYCN*) transgenic mouse provides an established model of high-risk neuroblastoma. In this genetically-engineered model, the *MYCN* gene, commonly amplified in human tumors, is the principal oncogenic driver and is highly expressed under the control of a tyrosine hydroxylase promoter to target tumor development to paraspinal or intra-abdominal sympathoadrenal tissues.[13] In addition to sharing *MYCN* overexpression, *TH-MYCN* tumors resemble human neuroblastomas histologically and genetically. [13–16] The model shows strain-dependent differences in phenotype, and, in the most widely utilized 129×1/SvJ background, mice homozygous for the *TH-MYCN* transgene (*TH-MYCN*^+/+^) have 100% lethal tumor penetrance.[14] Importantly, while each tumor arising in these mice shares *MYCN* overexpression as the initiating event, they have distinct cooperating mutations that promote transformation and undergo a unique immunoediting process in situ. *TH-MYCN* mice have an intact immune system, enabling the assessment of immune responses and immunotherapies. Importantly, unlike transplantable syngeneic tumor models that replicate tumors from a single tumor-derived cell line and implant them at an orthotopic or remote site, tumors in the *TH-MYCN* model initiate within autochthonous sympathoadrenal tissues, modeling interactions among neuroblasts, stromal cells and immune cells throughout the course of tumor initiation and progression. The tumor niche is important in influencing cancer progression as evidenced by syngeneic neuroblastoma transplant models in which subcutaneous implantation is associated with reduced tumor growth and fewer infiltrating tumor-suppressive immune cells compared with the same tumor cells injected at orthotopic intra-adrenal sites. [17,18]

Despite these advantages, an impediment to adoption of the highly penetrant 129×1/SvJ *TH-MYCN* model for studies into GD2-directed immunotherapies is the prevailing notion that these tumors lack sufficient surface GD2 to serve as an effective tumor antigen, primarily when investigated for syngeneic tumor transplantation studies. In the C57BL/6 background, preferred by many for immunologic studies, tumor penetrance for the *TH-MYCN* transgene is too low for efficient autochthonous modeling, and tumor-derived cell lines from this strain are found to express insufficient surface GD2 for therapeutic targeting unless they are manipulated pharmacologically or genetically. [19–20] Here, we demonstrate that neuroblastomas arising in 129×1/SvJ *TH-MYCN* mice express levels of GD2 nearly comparable to that present on human neuroblastomas. Importantly, not only is GD2 surface expression present on primary tumors in situ, but this antigen can be pharmacologically targeted using a GD2-directed monoclonal antibody, 14G2a, that is analogous to the human therapeutic antibody, dinutuximab. Therapy with 14G2a extends survival and provides durable complete tumor regression in a subset of treated mice. This survival benefit is accompanied by alterations in the TME, including selection for GD2-negative neuroblasts and alterations in the macrophage and MDSC populations. We further show that *TH-MYCN* neuroblasts explanted from *TH-MYCN* tumors rapidly lose GD2 expression while carried in standard tissue culture conditions, and that GD2 loss is associated with a transition of tumor cell lineage from an adrenergic to a mesenchymal state. Indeed, such adrenergic-mesenchymal lineage plasticity has been defined in human neuroblastomas with the mesenchymal state being correlated with therapy resistance and relapse.[21,22] Collectively, our work credentials the *TH-MYCN* model as a relevant platform for immunotherapeutic studies, including GD2-targeting approaches, and enables further studies into the regulation of mesenchymal state switch that renders tumor cells resistant to cytotoxic therapies and GD2-directed immunotherapies.

## METHODS

### Mice and treatments

129×1/SvJ-Tg *TH-MYCN* mice were originally obtained from Dr. William Weiss (University of California, San Francisco). [13] All mice were bred and housed in the animal facility at the Children’s Hospital of Philadelphia under approved IACUC protocols. Heterozygous *TH-MYCN*^+/−^ mice were bred and all studies performed on *TH-MYCN*^+/+^ offspring in which tumor-onset is fully penetrant. *MYCN* transgene genotype was determined from DNA isolated from a 1-cm tail snip by qPCR genotyping with the following primers:

Chr18F1 (5’-ACTAATTCTCCTCTCTCTGCCAGTATTTGC-3’),
Chr18R2 (5’-TGCCTTATCCAAAATATAAATGCCCAGCAG-3’),
and OUT1 (5’-TTGGCACACACAAATGTATATACACAATGG-3’), as described previously. [23]

Homozygous *TH-MYCN* mice were randomized to receive intraperitoneal injections of 100 μg of anti-GD2 antibody (14G2a; BioLegend), 100 μg of a murine isotype-matched antibody to control for Fc receptor binding (murine IgG2a/κ; Abcam), or phosphate-buffered saline (PBS), twice weekly starting between 14-17 days of age and continuing until day 100 of life. Mice were palpated for tumor presence and monitored for signs of morbidity that included poor mobility, hunching, lethargy, weight loss, limb paralysis, neurological changes, dermatitis, and rough hair coat. Mice exhibiting such signs were euthanized and underwent necropsy at “terminal” timepoints.

To assess the biological impact of 14G2a immunotherapy, an additional cohort of *TH-MYCN*^+/+^ mice (n=6) were treated with twice weekly 14G2a antibody beginning between day 14-17 of life and monitored as above. Regressing tumors were detected by abdominal palpation, typically between days 70-75, and harvested for characterization (called “midpoint tumors”).

*TH-MYCN*^+/+^ mice that survived to day 200 (alive without tumor symptoms >3 months from the end of therapy) were re-challenged with a single flank inoculation of HOM2 cells, a tumorigenic syngeneic *TH-MYCN*^+/+^ tumor-derived cell line (from Dr. Garrett Brodeur, Children’s Hospital of Philadelphia). Heterozygous *TH-MYCM*^+/−^ and wild-type (*TH-MYCN*^−/−^) 129×1/SvJ mice served as controls (n=4 each) and received a single flank injection on the same day using the same cell line preparation. Flanks were shaved and injected with 5×10^6^ cells resuspended in Matrigel (Corning Inc.) at a 1:1 ratio by volume. Mice did not receive any treatment after tumor inoculation. The proportion of mice with tumor engraftment, and time to engraftment were recorded. Tumors were procured for flow cytometric analysis, as below.

### TH-MFCN^+/+^-derived cell lines

Tumor-bearing mice were euthanized, and tumors removed and dissociated into small tissue fragments using the bottom of a sterile 3-ml syringe pistol in 5 ml of sterile PBS in a 6-cm petri dish. The cellular suspensions were filtered through a 40 μm sterile nylon mesh, washed multiple times, and resuspended in sterile TAC buffer (UltraPure Tris Hydrochloride; Invitrogen and Ammonium Chloride; Sigma). Cells were incubated in a 37°C water bath, spun, and resuspended in PBS. These steps were repeated until red blood cells were no longer visible. Pellets were then resuspended in complete IMDM (Iscove’s modified Dulbecco’s media) with 20% fetal bovine serum (FBS), 1% Penicillin-Streptomycin, 1% L-glutamine, 0.6% recombinant human insulin, human transferrin, and sodium selenite (ITS), and 0.25% gentamicin, transferred into a T75 flask via a 40 μm cell strainer and incubated at 37°C with 5% CO_2_. Approximately 60% of explanted tumors generate a cell line. Four *TH-MYCN*^+/+^ tumors were explanted to derive new cell lines (5144, 5150, 5195, and 5213) that were serially passaged and analyzed by flow cytometry analysis to assess GD2 expression and lineage defining markers over time. In addition, cryopreserved *TH-MYCN*^+/+^ tumor-derived cell lines that had been established previously and passaged > 3 months (3401, 3261, 3393, and 3392), and 3 isogenic *TH-MYCN*^+/+^ tumor cell lines obtained from the Brodeur laboratory (HOM2, G2 and G3B) were evaluated with flow cytometry for GD2 expression and lineage defining markers.

### Human neuroblastoma cell lines

Description of human neuroblastoma cell lines and culture conditions described in Supplemental Methods.

### Flow cytometry to assess GD2 expression and characterize tumor-infiltrating leukocytes

Tumor-bearing mice were euthanized at midpoint or terminal timepoints. Tumor fragments of 1-3mm^3^ were placed in RPMI Medium 1640 with L-glutamine (Gibco) and 10% heat-inactivated fetal bovine serum (FBS, Life Technologies) before undergoing tissue dissociation with a GentleMACS™ dissociator (Miltenyi Biotech) using the manufacturer’s recommended protocol with media including Collagenase Type IV (STEMCELL Technologies, Cambridge MA) and DNAse I (Sigma-Aldrich). After dissociation, suspensions were passed via a 70 μm filter, washed, and subjected to red blood cell lysis with ACK buffer (Gibco). Single cell samples were then frozen in fetal bovine serum with 10% dimethyl sulfoxide (Thermo Fisher Scientific) and stored at −80°C before staining for flow cytometry. Primary anti-mouse and anti-human antibodies used in experiments are listed in Supplemental Methods.

Cells were washed in PBS without serum and incubated with Zombie Aqua viability dye (BioLegend) according to manufacturer’s instructions and subsequently washed with FACS buffer (PBS supplemented with 2.5% fetal bovine serum; Thermo Fisher Scientific) before Fc blockade with TruStain FcX (BioLegend) and subsequent surface marker staining. Cells were analyzed on a FACSVerse™ flow cytometer (BD Bioscience). Data analysis of flow data and gating strategy are discussed in Supplemental Methods.

### Detection of lineage-defining proteins

Description of immunoblotting and immunocytology are described in supplemental methods.

### Statistical analyses

Statistical analysis was performed using Prism software (GraphPad). Survival curves were compared using the method of Kaplan-Meier with a log-rank test for significance. For comparison of groups in flow cytometry analysis, groups were compared using unpaired two-tailed Student’s *t-*tests. For all, significance was set as p<0.05.

## RESULTS

### GD2 is highly expressed on tumor cells within *TH-MYCN* neuroblastomas in situ but is lost during propagation as tumor-derived cell lines

End-stage intra-abdominal *TH-MYCN*^+/+^ mouse primary tumors were harvested at the time of sacrifice for symptomatic progression and examined for the expression of GD2 on neuroblasts. All tumors expressed GD2 on the surface of the majority of tumor cells (n=7; median % positive: 86%, range 56-96%; **Fig.1A**). We next assessed GD2 expression on cell lines previously established and passaged from tumors arising in our *TH-MYCN* colony (n=4). Despite robust GD2 expression on primary tumor cells, tumor-derived cell lines uniformly had <5% GD2 expression (**Fig.1A**) though they remained tumorigenic and expressed neuroblast markers. To assess for temporal variability in surface GD2 levels, we passaged these *TH-MYCN* cell lines weekly yet GD2 expression remained consistently low. We compared this with GD2 expression on human neuroblastoma cell lines (n=10). These latter cell lines retained stable and high GD2 expression despite having been passaged extensively in tissue culture, with the exception of one outlier with lower frequencies of GD2^+^cells (**Fig.1A**).

**Figure 1.**
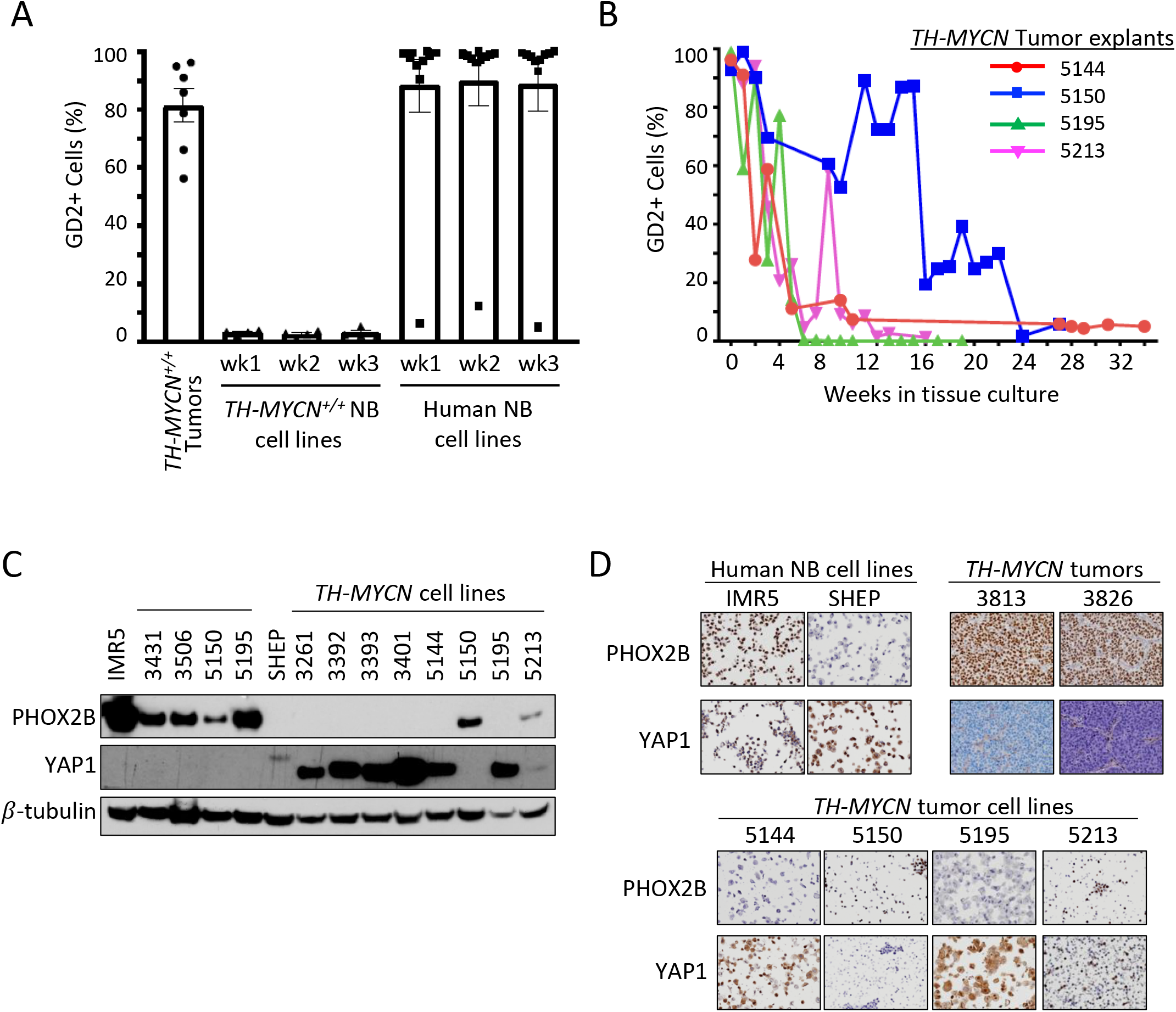
GD2 and lineage marker expression in *TH-MYCN* tumors, primary explants and cell lines, and human neuroblastoma cell lines. (A) *TH-MYCN*^+/+^ neuroblastomas (NB), previously established and propagated *TH-MYCN*^+/+^ and human NB cell lines were examined for surface GD2 expression by flow cytometry in single, live, CD45^-^ cells (and NCAM1^+^ and CD56^+^ for mouse and human NBs, respectively). Cell lines were also passaged for 3 weeks to assess GD2 stability. (B) Fresh *TH-MYCN*^+/+^ tumors were explanted and carried in tissue culture to monitor the GD2^+^ proportion of tumor cells over time ex vivo. (C) Immunoblot detection of a principal adrenergic marker, Phox2b, principal mesenchymal marker, Yap1, and β-tubulin loading control. IMR5 and SHEP are neuroblastoma cell lines with a predominantly adrenergic or mesenchymal lineage, respectively. *TH-MYCN*^+/+^ primary tumors are predominantly adrenergic marker expressing. *TH-MYCN*^+/+^ tumor-derived cell lines are largely mesenchymal marker expressing when propagated ex vivo (>3 months, 3000-series cell lines) whereas marker expression is heterogeneous during earlier passage ex vivo (5000-series cell lines). (B) Immunocytochemistry and immunohistochemistry to detect Phox2b and Yap1 from FFPE cell line pellets or primary tumors are shown, concordant with immunoblot results.

To reconcile these findings, we explanted four *TH-MYCN* primary tumors into tissue culture and serially examined the expression of GD2 over time from explant through outgrowth of a cell line. Nearly all CD45^−^/NCAM1^+^ live neuroblasts expressed high levels of GD2 at the time of initiation of the culture (with a median of 95% GD2^+^/NCAM1^+^/CD45^−^ live cells, with a range of 93-99%). However, the proportion of such tumor cells expressing GD2 subsequently decreased over time (**Fig.1B**). The kinetics of GD2 loss varied but all reached a stable low or absent level between 6-24 weeks of propagation. Therefore, while human neuroblastoma cells maintain GD2 expression while propagated ex vivo, *TH-MYCN-derived* cell lines lose surface GD2 expression soon after they are explanted to tissue culture conditions despite the absence of selective pressure to do so.

### *TH-MYCN* tumors possess an adrenergic-predominant state, whereas *TH-MYCN* tumor-derived cell lines possess a mesenchymal-predominant state

Human neuroblasts and cell lines have lineage plasticity and can trans-differentiate between an adrenergic (sympathetic) and mesenchymal (neural-crest-like) state. [21,22] Since these epigenetic phenotypes associate with distinct transcriptional outputs, we determined the lineage state of *TH-MYCN* tumor cells using a consensus adrenergic marker, Phox2b, and a consensus mesenchymal marker, Yap1 by immunoblot, immunocytology, and/or immunohistochemistry. [21,22] All *TH-MYCN* primary tumors expressed abundant Phox2b but absent Yap1, supporting an adrenergic-predominant state (**Fig.1C-D**) in association with their high GD2 expression. In contrast, all established *TH-MYCN* tumor-derived cell lines expressed abundant Yap1 and absent Phox2b, consistent with a mesenchymal-predominant state (**Fig.1C-D**; **Fig.S1A**) in association with their low GD2 expression. We next studied *TH-MYCN* tumor explants at intermediate time-points in tissue culture. The nascent *TH-MYCN* cell lines 5144 and 5195 both lost surface GD2 expression rapidly (**Fig.1B**) and expressed mesenchymal markers even at early time-points (**Fig.1C**). However, the two *TH-MYCN* cell lines with prolonged retention of GD2 expression were found to retain an adrenergic expression pattern (5150) or express both markers (5213) when assessed at intermediate time-points. Overall, there was striking concordance between GD2 expression and adrenergic lineage in the *TH-MYCN* models. These data support a lineage switch from adrenergic to mesenchymal state when murine neuroblasts are cultured ex vivo leading to loss of GD2 expression, likely due to loss of adrenergic transcriptional programs. To assess whether 5213 cells included tumor cells that co-expressed Phox2b and Yap1 or an admixture of both lineage states, we studied cell pellets by immunocytology, which showed a population of Phox2b expressing cells admixed with a population of Yap1 expressing cells (**Fig.1D**). We next determined whether the tissue-culture adopted mesenchymal state could readily revert to the adrenergic state in situ, as human neuroblastoma cell lines do when re-implanted as tumor xenografts. [21] We allowed *TH-MYCN* cell lines to establish as xenografts in the flank of syngeneic mice and then used immunohistochemistry to detect Phox2b and Yap1, demonstrating that they retained their mesenchymal status (**Fig.S1B–S1C**).

### Treatment of *TH-MYCN* tumor-bearing mice with an anti-GD2 antibody induces tumor regression and extends survival

Since *TH-MYCN* neuroblasts retain high levels of GD2 expression in situ despite losing GD2 expression in tissue culture, we sought to determine the effect of treatment with an anti-GD2 antibody. *TH-MYCN* mice (n=15 per treatment arm) were randomly assigned to therapy with 14G2a (a murine anti-GD2 antibody), an isotype-matched control antibody, or PBS (**Fig.2A**). Mice received their assigned treatment twice weekly from day of life 14-17 (the time of tumor initiation based on histologic audits) until they were sacrificed for symptoms of tumor progression or until day 100. [24] All mice treated with an isotype-control antibody or PBS had progressive tumor growth and were euthanized while still receiving therapy (median survival time of 46 and 49 days, respectively: **Fig.2B**). In contrast, mice treated with 14G2a antibody had significantly extended survival (median survival time of 81 days; p<0.001 compared with either control arm) with 7 mice completing assigned treatment through day 100, beyond which they continued to be observed for signs of tumor progression. 14G2a monotherapy led to objective tumor regression in 6 of 15 mice (40%), and 4 of these mice (27%) had complete regression to palpation that was maintained until day of life 200. All four of these mice were subsequently re-challenged with a flank inoculation of a syngeneic *TH-MYCN* tumor cell line (details below) and at the time of necropsy had no residual tumor at their primary autochthonous tumor site.

**Figure 2.**
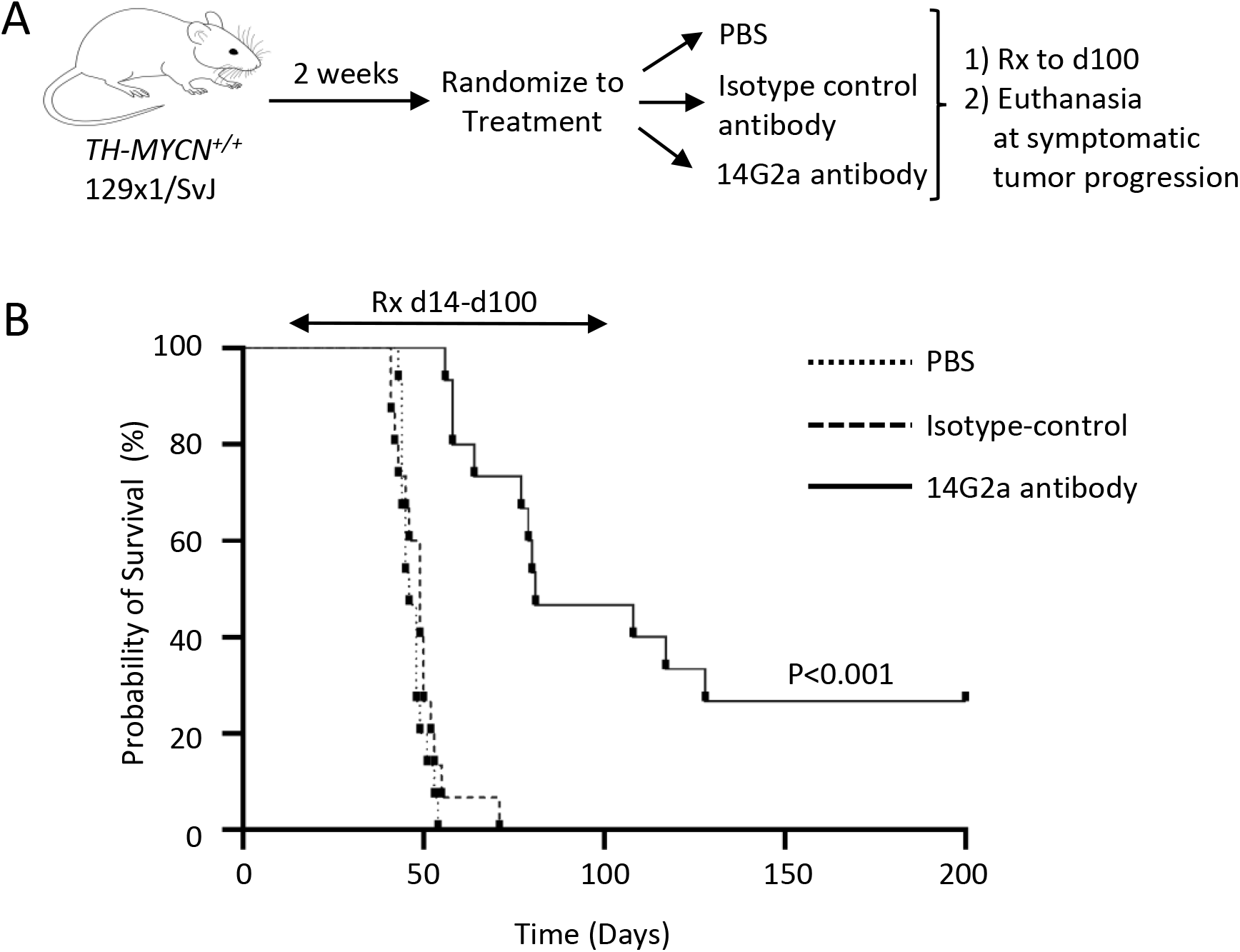
Experimental Summary and survival of *TH-MYCN*^+/+^ mice. (A)Transgenic *TH-MYCN*^+/+^ mice were enrolled into 3 treatment groups: PBS, 14G2a, and isotype. Mice were treated with respective intraperitoneal (IP) injections twice weekly until death or significant morbidity, and then had tumors isolated in terminal surgeries. (B) Kaplan-Meier survival curve shown by treatment group (note: 4 mice living at day 200 in the 14G2a-treated group were censored for rechallenge with flank implantation of a *TH-MYCN-derived* neuroblastoma cell line). Log-rank (Mantel-Cox) test was used to compare survival curves with significance set at p<0.05.

We assessed the stability of GD2 expression on tumors over time in situ by correlating the proportion of GD2^+^ tumor cells with days of survival (age). No correlation was found with age (time of treatment) for mice treated with PBS or isotype-control antibody, although the range evaluable was modest due to early tumor lethality. The 14G2a-treated cohort included mice treated in the survival study with tumors harvested at the time of terminal tumor progression (terminal), and additional mice whose tumors were harvested at the time of GD2 antibody-mediated tumor regression (midpoint). We again saw no correlation between the proportion of GD2^+^ tumor cells with survival time within the GD2-antibody treated group (**Fig.3C**). However, comparing GD2 expression among tumors harvested at the time of sacrifice (terminal tumors) from different treatment groups showed significantly reduced GD2 on tumors from 14G2a-treated mice, with a reduced proportion of GD2^+^ cells and reduced GD2 mean fluorescence intensity (MFI) compared with PBS treated tumors (p=0.03 for each), and non-significant reductions in both compared with isotype-control antibody treated tumors (p=0.14 and 0.13, respectively; **Fig.3A-B**). 14G2a-treated tumors harvested at the time of regression (midpoint) also had a lower proportion of GD2^+^ cells than isotype-antibody or PBS treated terminal tumors (p=0.06 trend, and p<0.01, respectively), and reduced MFI compared with PBS treated tumors (p=0.02). Notably, GD2 MFI was lower in 14G2a-treated midpoint tumors compared with terminal tumors (p=0.04). This may reflect the impact of maximal selective pressure against GD2 expression at the time of tumor regression (midpoint), and that more than a quarter of 14G2a-treated mice survived beyond day 100 and had stopped 14G2a treatment prior to the time of sacrifice.

**Figure 3.**
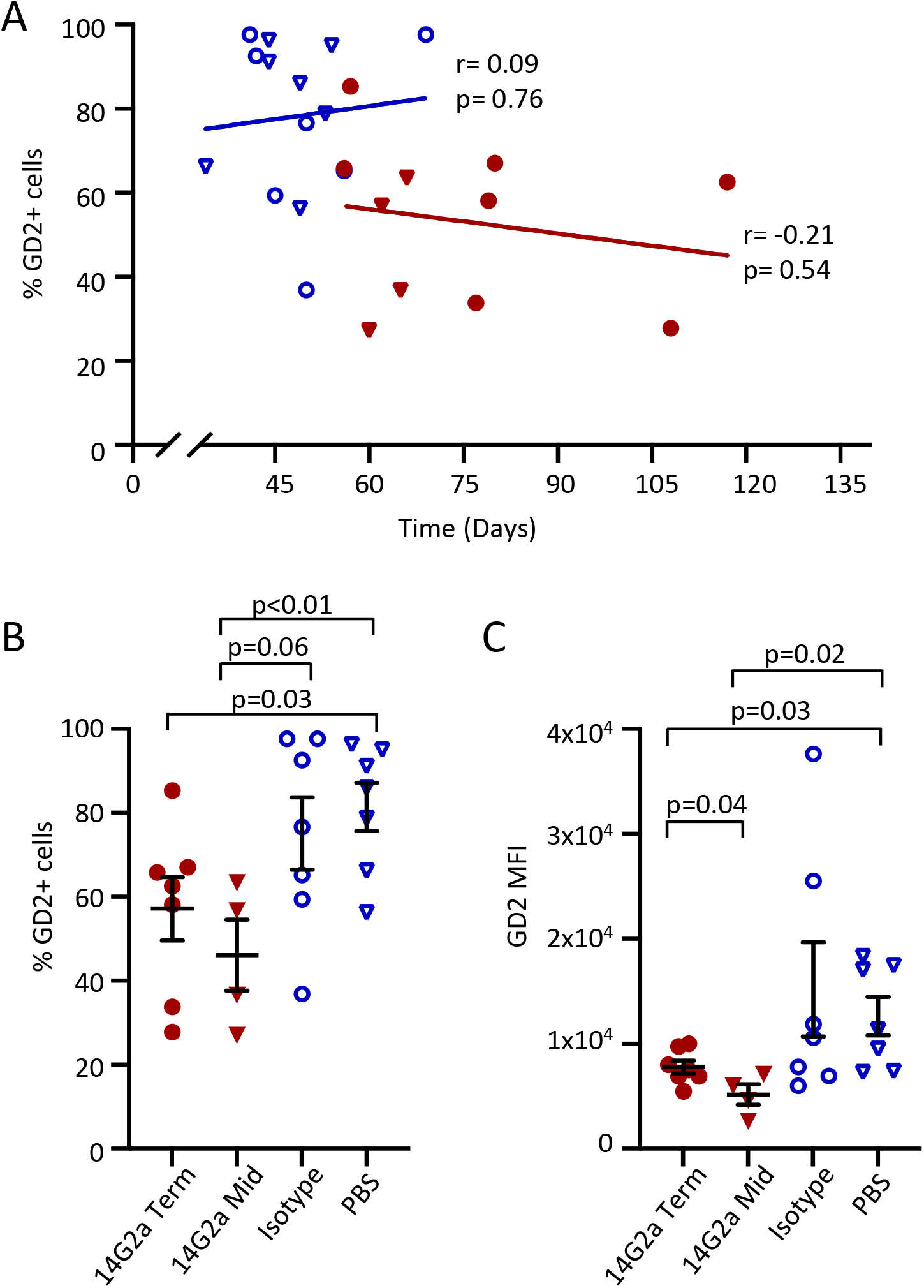
GD2 Expression in *TH-MYCN*^+/+^ neuroblasts with 14G2a treatment and correlation with survival. Neuroblastoma tumors isolated from *TH-MYCN*^+/+^ mice in 14G2a terminal (Term) [red circles], 14G2a midpoint (Mid) [red triangles], Isotype control [blue open circles], and PBS [blue open triangles] treatment groups were processed into single cell suspensions and immunostained. Single, CD45^−^, live cells were analyzed for cell surface expression of GD2. (A) Simple linear regression of survival in days and percent GD2^+^ cells in 14G2a-treated (GD2 Term and GD2 Mid) and non-14G2a-treated (Isotype and PBS) groups. (B and C) The percent of GD2^+^ cells and GD2 mean fluorescence intensity (MFI) in tumors from *TH-MYCN* mice is shown by treatment group. Significance determined by two-tailed Student’s t-test in A&B, and simple linear regression in C with significance set as p=0.05 (trend p<0.10).

### *TH-MYCN* mice achieving durable tumor control following anti-GD2 therapy are not protected against re-challenge with syngeneic tumor

Four 14G2a-treated *TH-MYCN* mice appeared healthy with no palpable tumor at day 200 (100 days from stopping therapy). To assess whether a memory T cell response had developed in these mice in response to neo-epitopes uncovered as a result of the antibody therapy or other immune changes evoked, we re-challenged them with a subcutaneous injection of a tumorigenic *TH-MYCN-derived* cell line (HOM2 cells). Tumor-free *TH-MYCN*^+/−^ (n=4) and *TH-MYCN*^−/−^ (n=4) mice were concurrently challenged. Tumorigenicity was 100% across all mice, with firm growing flank tumors at a median of 11.5 and 9 days, respectively, for *TH-MYCN*^+/−^ and *TH-MYCN*^−/-^ (wild-type) control mice, and 15 days in re-challenged *TH-MYCN* mice whose original autochthonous tumors regressed with prior anti-GD2 therapy. There were not statistically significant differences in time to engraftment among groups. Of note, HOM2 *TH-MYCN* tumor cells express mesenchymal markers as do their established xenografted tumors (**Fig.S1C**).

### Characteristics of tumor-infiltrating immune cells in *TH-MYCN* tumors

*TH-MYCN* tumors recruit an immunosuppressive microenvironment similar to human neuroblastomas, with a reduction in cytolytic T cells and increase in TAMs, dendritic cells, and MDSCs. [25–27] To determine the impact 14G2a antibody therapy had on the TME, we first examined the overall frequencies of intra-tumoral leukocytes in terminal tumors. Conventional T cell, invariant natural killer T (iNKT) cell, gamma-delta (γδ) T cell, dendritic cell and NK cell frequencies were unchanged in 14G2a-treated tumors when compared with isotype-control antibody or PBS-treated tumors, respectively. However, TAMs, monocytic-MDSCs (M-MDSC) and granulocytic-MDSCs (G-MDSC) were all present at reduced frequencies in 14G2a-treated tumors compared with PBS-treated tumors, reflecting a less immunosuppressive TME (**Fig.4**).

**Figure 4:**
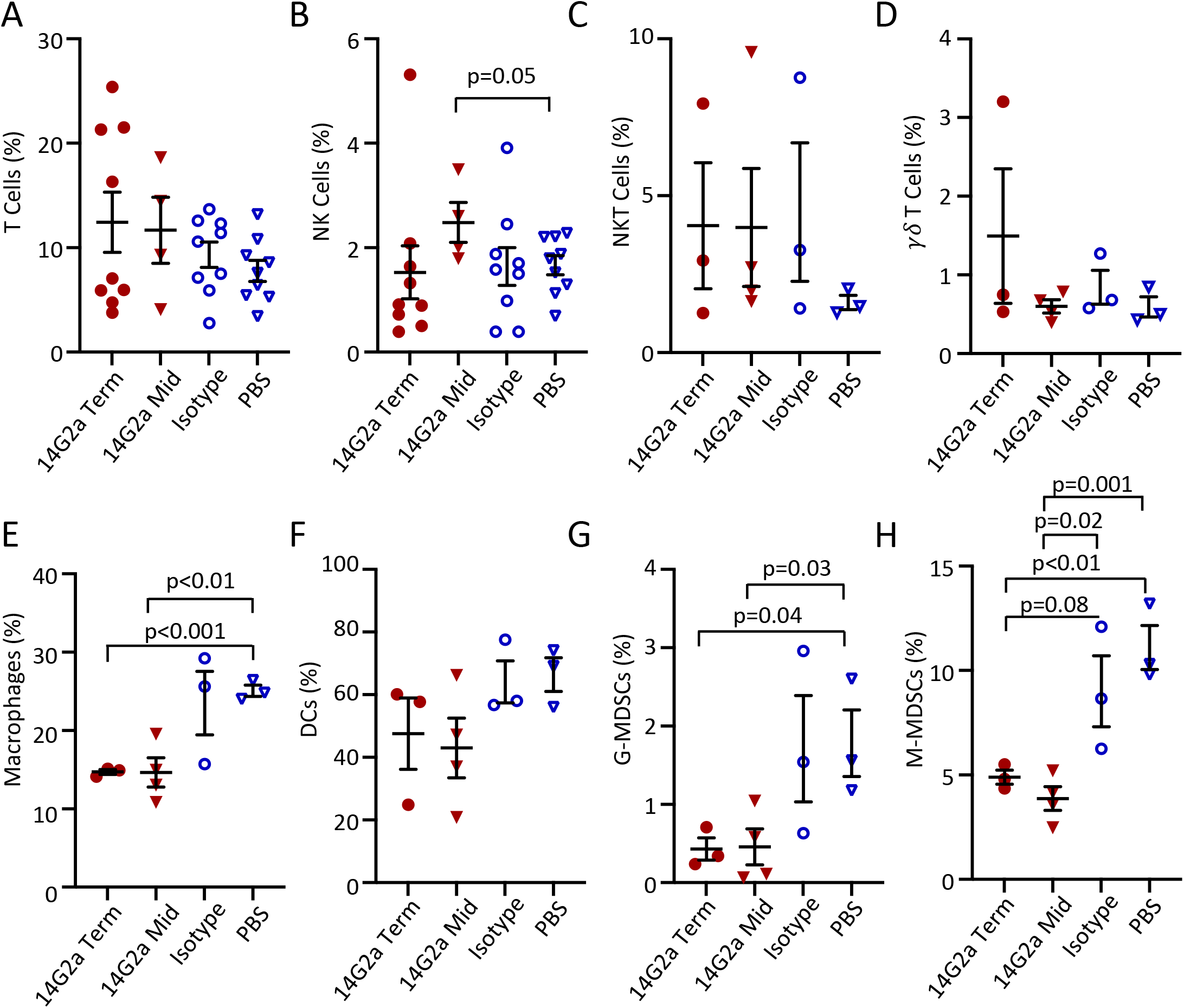
Immune Cells in the *TH-MYCN^+/+^Tumor* Microenvironment. *TH-MYCN*^+/+^ neuroblastoma tumors were isolated from 14G2a terminal (Term), 14G2a midpoint (Mid), isotype control, and PBS treatment groups. Single cell suspensions of tumors were immunostained with fluorescent antibodies specific for markers of natural killer (NK), natural killer T (NKT), gamma delta (γδ) T, and T cells in addition to markers for macrophage, dendritic cells (DCs), granulocytic myeloid-derived suppressor cells (G-MDSCs) and monocytic myeloid-derived suppressor cells (M-MDSCs). (A-H) Specified cell frequencies are reported as a percentage of single, live, CD45^+^ parent populations, by treatment group. Groups were compared with two-tailed Student’s t-test with significance set as p<0.05 (trend p<0.10).

Several 14G2a-treated mice were removed from therapy before being sacrificed for tumor progression, so we sought additional immune changes in midpoint tumors at the time of maximal immunotherapy impact. We confirmed the reduction in TAMs, M-MDSCs, and G-MDSCs but also identified an increase in NK cells within the tumors (**Fig.4B**). Since NK cells are postulated to be major effectors of the therapeutic response induced by GD2-directed antibody therapy, we examined their pattern of activating and inhibitory receptors and found a significant decrease in the frequencies of intratumoral NK cells expressing the activating Ly49H receptor and a trend towards greater frequencies of NK cells expressing inhibitory Ly49A and/or TGFβ receptor type 1 (TGFβR1) in 14G2a-treated tumors (**Fig.5A-C**). [28] There were no differences in the proportions of NK cells expressing NKG2D, NKG2A, or interleukin-15 receptor-α (IL-15Rα) among treatment groups at terminal timepoints. Finally, iNKT cell surface expression of CD107a was also noted to be higher in 14G2a relative to the isotype-control group (p=0.04). Overall, while reductions in TAM and MDSC frequencies and concomitant increases in NK cell proportions suggest that the balance of intratumoral immunosuppressive to effector cells may favor tumor control, surrogates of NK cell function such as activation and inhibitory receptors are inversely altered, suggesting that the effector function of tumor-infiltrating NK cells may be diminished, perhaps via exposure to TGFβ. Conversely, iNKT cells from 14G2a-treated tumors appear to have upregulated surrogates of cytotoxic function (CD107a expression), but their frequencies do not appear to be altered with treatment.

**Figure 5:**
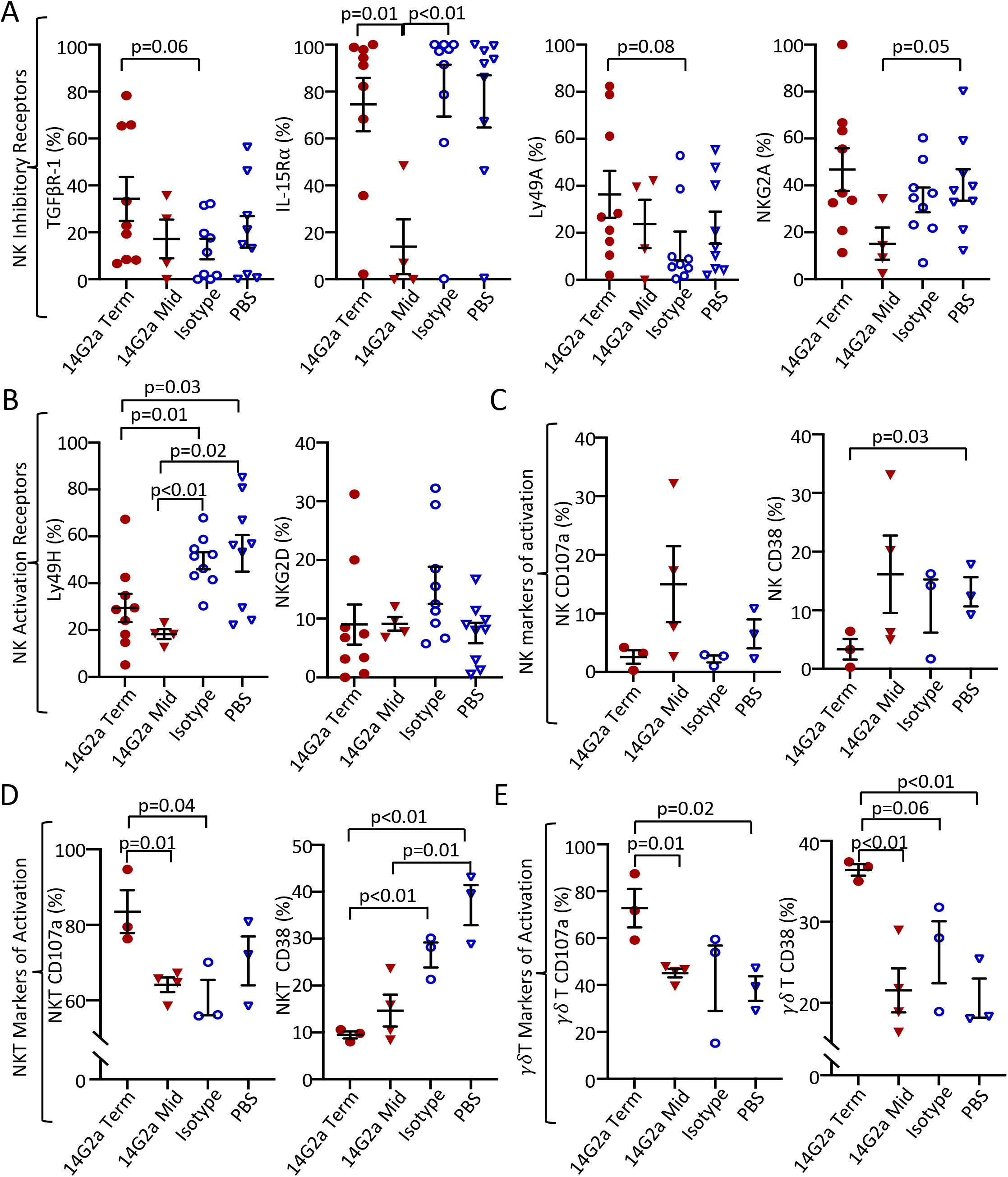
NK, NKT, and γδ T Cell surface markers in the *TH-MYCN*^+/+^ TME. Cell surface protein expression from *TH-MYCN*^+/+^ neuroblastoma-derived natural killer (NK), invariant natural killer T (NKT), and gamma delta (γδ) T Cells from 14G2a terminal (GD2 Term), 14G2a midpoint (GD2 Mid), Isotype, and PBS groups. (A) NK Inhibitory receptor expression including transforming growth factor beta receptor-1 (TGFBR-1), interleukin-15 receptor (IL-15R), Ly49A, and NKG2A. (B) NK Activation receptor expression of Ly49H and NKG2D. (C-E) Expression of markers of cellular activation CD38 and CD107a are shown for NK, NKT, and γδT cells. Two-tailed Student’s t-test performed to compare groups with significance set as p<0.05 (trend p<0.10).

To further probe whether the changes noted in intratumoral NK cells reflected alterations in the susceptibility of neuroblasts to immune control, we examined the tumor cell expression of ligands that regulate NK cell reactivity to putative targets, including NK cell activating ligands retinoic acid early-inducible protein 1-gamma (Rae-1γ) and UL16 binding protein 1 (ULBP-1), and NK cell inhibiting ligands major histocompatibility class I (MHC-I) and latency-associated peptide (LAP; **Fig.6A-D**). We found that a higher proportion of 14G2a-treated neuroblasts expressed Rae-1γ relative to PBS-control (trend, p=0.07), while a lower proportion expressed ULBP-1 (p<0.01) and MHC-I (trend, p=0.06). ULBP-1 expression in 14G2a-treated tumors was also decreased relative to isotype-treated tumors (p=0.04). The frequencies of LAP-expressing neuroblasts were lower in 14G2a-treated versus PBS-treated mice (p=0.04). As NK cells are activated via binding of NKG2D to ULBP-, and CD8^+^ T cells recognize tumors antigens on MHC-I, the decrease in ULBP-1 and MHC-I in 14G2a-treated tumors may reflect mechanisms for neuroblast immune evasion under increased immune pressure with 14G2a treatment. [29–30] Collectively, these data highlight the complexities of interactions and alterations in the *TH-MYCN* neuroblastoma TME associated with anti-GD2 treatments.

**Figure 6.**
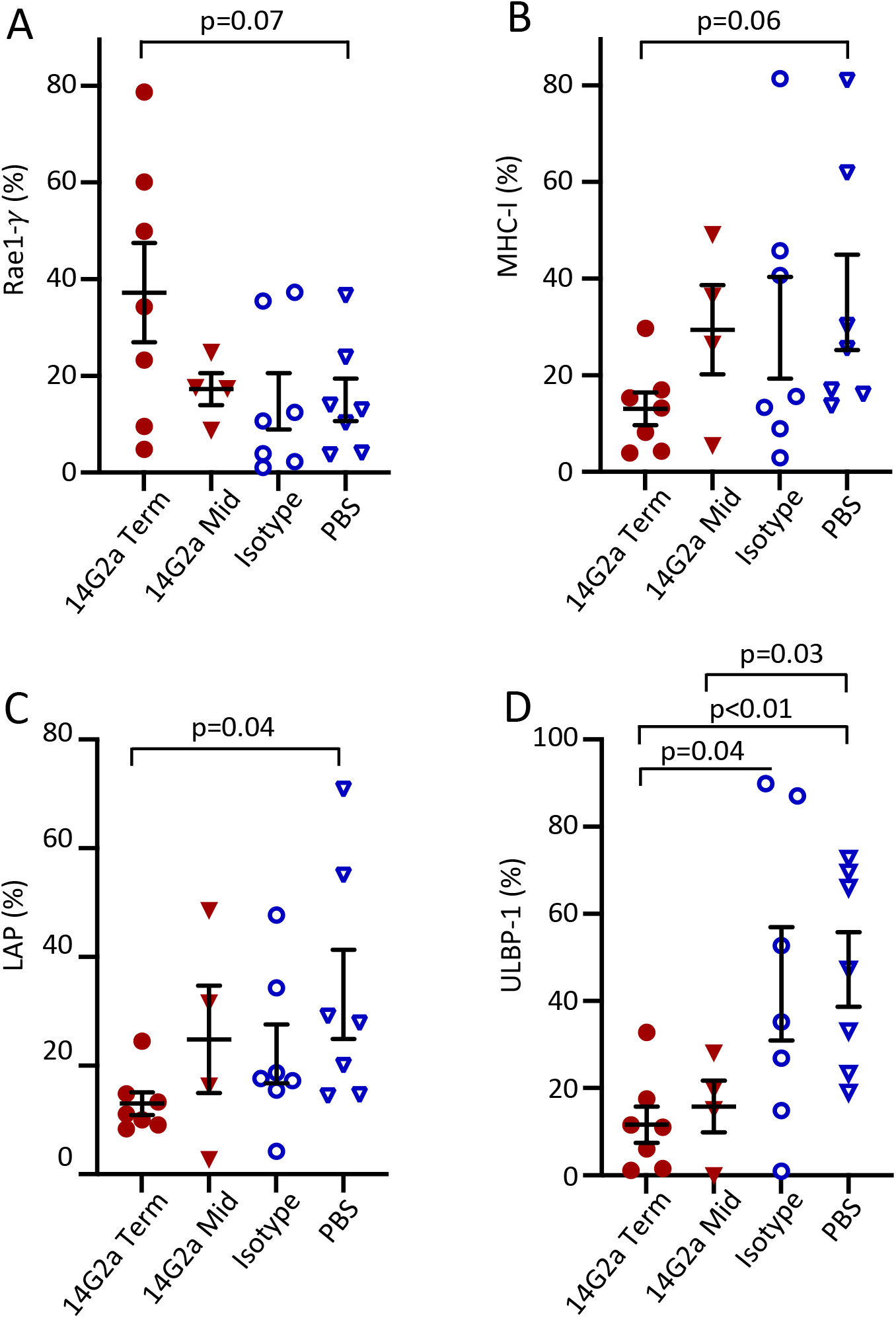
Surface expression of NK Ligands on *TH-MYCN*^+/+^ Neuroblasts. Tumors from 14G2a terminal (Term), 14G2a midpoint (Mid), Isotype control, and PBS-treated *TH-MYCN*^+/+^ mice were isolated. Single cell suspensions were immunostained and CD45-, GD2^+^, live cells were analyzed for cell surface expression of specified NK Ligands. (A-D) Retinoic acid early-inducible protein 1-γ(Rae1-γ), Major Histocompatibility Complex Class I (MHC-I), Latency Associated Peptide (LAP) and UL16 Binding Protein-1 (ULBP-1) expression on live, CD45^−^, GD2^+^ neuroblasts is shown. Bars show the mean and standard error of the mean. Groups were compared with two-tailed Student’s t-test with significance set as p<0.05 (trend p<0.10).

## DISCUSSION

The use of models of cancer that faithfully recapitulate human disease has enabled the discovery and preclinical prioritization of novel therapies. Here we credential the *TH-MYCN* transgenic mouse for neuroblastoma immunotherapy studies. *TH-MYCNs* use the principal oncogenic driver, *MYCN*, to initiate tumors at autochthonous sympathoadrenal sites whose distinct cooperating mutations influence the immunoediting process. This in situ immunoediting results in the recapitulation of the genomic, microenvironmental, and immunologic heterogeneity that defines cancer. As such, the *TH-MYCN* model represents distinct advantages over syngeneic transplant models for investigating the activity of immunotherapies such as dinutuximab. Although the model is well established, it has been underutilized as a platform to test GD2-directed therapies due to the pervasive impression that *TH-MYCN* tumor cells express insufficient surface GD2 to be targeted. Here we show that tumors in situ express GD2 levels at high frequencies, and that this expression is therapeutically actionable. We observed a profound survival benefit with 14G2a therapy with near doubling of median survival time, 40% of mice with objective tumor regressions, and 30% with durable complete responses beyond 200 days. Similar results using *TH-MYCN*^+/−^ mice were recently shown, along with a synergistic effect of adding chemotherapy to GD2-directed antibody, analogous to the findings in children with relapsed neuroblastoma in the COG ANBL1221 chemoimmunotherapy trial. [8,17]

Moreover, we found a reduced proportion of neuroblasts from 14G2a-treated mice expressed GD2 with decreased MFI when compared with PBS-treated tumors at both the midpoint and terminal time points, suggesting a selective pressure was exerted by 14G2a treatment. Recent work has suggested that GD2 serves as a macrophage checkpoint or “don’t eat me” signal between neuroblasts and TAM. [31] The interplay of GD2-ligation with 14G2a and macrophage and MDSC clearance may therefore be related to immunologic activation via FcR binding, via cell-cell interaction changes mediated by blockade of GD2-macrophage signaling, or by complement binding. Notably, 7 of 15 14G2a mice lived without morbidity beyond cessation of antibody administration at day 100, while none of the mice in the control groups survived beyond day 70. This finding prompted the hypothesis that mice with prolonged survival had gained immunologic memory due to epitope-spreading from the anti-GD2 antibody therapy. This was tested in a re-challenge study in the 4 mice that lived past day 200 without signs of morbidity. However, there was no decrease in tumorigenicity or time to tumor establishment between these mice and untreated controls, suggesting a lack of long-term immunity with the use of 14G2a in this model. However, this finding may be confounded by the switch in lineage state in the cell lines used for re-challenge. It remains possible that long-term adaptive immunity developed but was directed at adrenergic state antigens that are lost in the mesenchymal state.

We also show that established *TH-MYCN* tumor-derived cell lines have markedly reduced GD2 expression in the absence of prior exposure to an anti-GD2 immunotherapy, confirming the findings of others. [18,20] Importantly, *TH-MYCN* neuroblasts lose GD2 during the process of explanting the primary tumor and through propagation as a cell line, such that serial syngeneic transplant of such tumor xenografts is unlikely to be useful for GD2-directed studies, as shown previously. [18,20] The majority of GD2 expression was lost in <8 weeks of propagation, suggesting the in situ environment provided a stimulus for GD2 expression, or that tissue culture conditions induced its loss. It has previously been shown that neuroblastomas exist in distinct epigenetic states that can interconvert. [22] Indeed, we show that a change in lineage state with conversion from a predominantly adrenergic state to a mesenchymal state accompanies loss of GD2 from the tumor cell surface. This change in state has been genetically associated with NOTCH-family transcription factor signaling, loss of *ARID1A* expression, and with TNF-α and epidermal growth factor exposure in vitro. [32–34] It has previously been suggested that GD2 is expressed more highly at cell-cell junctions and it is possible that disruption of these contacts and removal from the sympatho-adrenal location contribute to GD2 loss in tissue culture. [35] Notably, expression of GD2 synthases, in particular the *B4GALNT1* enzyme that catalyzes the final step in synthesis, is confined to sympathoblasts (adrenergic) in single-cell RNAseq studies. [36] Plausibly, GD2 expression is downstream of the adrenergic state transcriptional output and is lost as a consequence of lineage change to the mesenchymal state. Our finding that *TH-MYCN-* derived tumors cells can assume a mesenchymal state both in culture and in re-implanted tumors may prove important. Emerging evidence suggests that adrenergic and mesenchymal states have profound differences in inflammatory signaling and immune susceptibility, and an immunocompetent model to study these states is lacking. [37]

Dinutuximab is purported to exert its antitumor effect via antibody-dependent cellular cytotoxicity (ADCC) as well as complement-dependent cytotoxicity. [7] In our evaluation of the *TH-MYCN* TME, we did not observe differences in intratumoral NK, T, and iNKT cell frequencies between 14G2a and isotype control groups at terminal timepoints, but did observe an increase in NK cells at the time of response to 14G2a (midpoint tumors) compared with the PBS control. NK cells are known to transactivate iNKT cells and we did observe an increase in CD107a expression in 14G2a-treated intratumoral iNKT cells. [38] These findings suggest that that anti-GD2 antibody may result in downstream iNKT cell degranulation. Additionally, it has been shown that activated iNKT cells reshape the immunosuppressive TME through recognition and lysis or reprogramming of TAMs and MDSCs. [39] It was therefore interesting to find that macrophage, M-MDSC, and G-MDSC populations were reduced in 14G2a-treated groups relative to PBS and isotype controls. As TAMs and MDSCs are known to provide critical support for NB tumor cell growth and are enriched in metastases, the reduced intratumoral frequency of these cells in the 14G2a-treatment group may represent a beneficial effect of 14G2a contributing to the prolonged survival seen in *TH-MYCN* mice. [27,40–42]

Upon further examination of the intratumoral NK cells, we observed a trend towards a larger percentage of these lymphocytes bearing the inhibitory receptors TGFβR-1 and Ly49A and a lower percentage expressing the activating receptor Ly49H in the 14G2a relative to the isotype control antibody cohort. We speculate that this finding indicates that the mice who eventually succumb to disease are ones in whom the 14G2a-induced immune pressure has diminished. This altered NK receptor profile following anti-GD2 antibody treatment provides potential opportunities for enhancing 14G2a-mediated efficacy. For instance, binding of TGFβ to the increased TGFβ-R1 could diminish NK cell gamma interferon-γ (IFN-γ) production and ADCC, thereby contributing to 14G2a resistance. [43] Use of TGFβ-R1 blockade with galunisertib has been used in vitro and in immunocompromised mice with improved efficacy of GD2-directed antibody therapy, but not in immunocompetent *TH-MYCN* mice. [44]

We also show that a lower frequency of *TH-MYCN* neuroblasts from 14G2a-treated mice express the NK cell activating ligand ULBP-1 than those from isotype-treated control mice. Conversely, a larger proportion of neuroblasts expressed Rae-1γ in mice treated with 14G2a relative to the isotype control antibody, although this did not reach statistical significance. Percentages of neuroblasts expressing LAP and MHC-I were decreased in 14G2a-treated terminal tumors relative to PBS-treated terminal tumors (p=0.04 for LAP expression, p=0.06 for MHC-I expression). MHC-I downregulation has previously been described as a mechanism for immune evasion and this finding in 14G2a-treated tumors may represent a response to increased immunologic pressure. [45]

The *TH-MYCN* mouse model offers many advantages for the study of immune:tumor interactions, acknowledging there are certain limitations including the low frequency of macrometastases. [13,16,17,46,47] Anti-GD2 therapies against neuroblastoma have been studied in alternative pre-clinical mouse models including those using orthotopic tumor injections of either human cell lines or patient-derived xenografts into immunodeficient mice.[16,48,49] These studies showed promising results regarding the augmented cytotoxic effect of anti-GD2 immunotherapy in combination with either soluble IL-15/IL-15Rα and GM-CSF or *ex vivo* activated human NK cells following tumor resection, but are limited by their immunodeficiency and altered TME. [48, 49] A combination of pre-clinical mouse models will likely serve as the ideal approach to study and evaluate the GD2-directed immunotherapy in neuroblastoma.

Collectively, these findings support the *TH-MYCN* immunocompetent mouse model’s utility for pre-clinical assessments of the efficacy of immunotherapies, including combination approaches that may synergize with GD2-targeted therapies. We also demonstrate the usefulness of the model for evaluating TME changes which can inform future therapeutic approaches. Such pre-clinical assessments have potential for expediting the identification of new treatment options for high risk neuroblastoma.

## Supporting information

GD2SuppMethodsFile

## DECLARATIONS

### Ethics approval and consent to participate

All animal studies were carried out under a protocol approved by IACUC at the Children’s Hospital of Philadelphia.

### Consent for publication

The authors give consent for publication of referenced data. There are no identifiable details or human subject data included in this manuscript.

### Availability of data and material

All data that support the findings of this study are available from the corresponding authors upon reasonable requests.

### Author Contributions

KOM and SK designed, performed experiments, data analysis, created figures and initial draft of the manuscript; FG designed, performed experiments, and data analysis; AW, CB, PK, RV, and AV provided technical expertise, performed experiments, and data analysis. MDH and HB conceived and supervised the study, were involved in the design and evaluation of experiments, critically reviewed all drafts of the manuscript. All authors contributed to the writing of the manuscript, critically reviewed, and approved the submitted version

## Acknowledgements

We would like to thank Dr. Garrett M. Brodeur (Division of Oncology, Children’s Hospital of Philadelphia) for kindly sharing the HOM2, G2 and G3B cell lines.

## LIST OF ABBREVIATIONS

|  |  |
| --- | --- |
| ADCC | Antibody-dependent cellular cytotoxicity |
| DCs | Dendritic Cells |
| FBS | Fetal bovine serum |
| $\gamma\delta$ T cell | Gamma delta T cell |
| G-MDSC | Granulocytic myeloid-derived suppressor cells |
| IFN- $\gamma$ | Interferon- Gamma |
| IMDM | Iscove's modified Dulbecco's Media |
| iNKT | Invariant natural killer T cells |
| LAP | Latency-associated peptide |
| MFI | Mean fluorescence intensity |
| MHC | Major histocompatibility complex |
| MDSC | Myeloid-derived suppressor cells |
| M-MDSC | Monocytic myeloid-derived suppressor cells |
| NK Cell | Natural killer cell |
| NCAM1 | Neural cell adhesion molecule-1 |
| PBS | Phosphate-buffered saline |
| TAM | Tumor associated macrophage |
| TGFβR1 | TGFβ receptor type 1 |
| TH-MYCN | Tyrosine Hydroxylase-MYCN |
| TME | Tumor microenvironment |
| ULBP-1 | UL16 binding protein 1 |

---

**Supplemental Figure 1.**
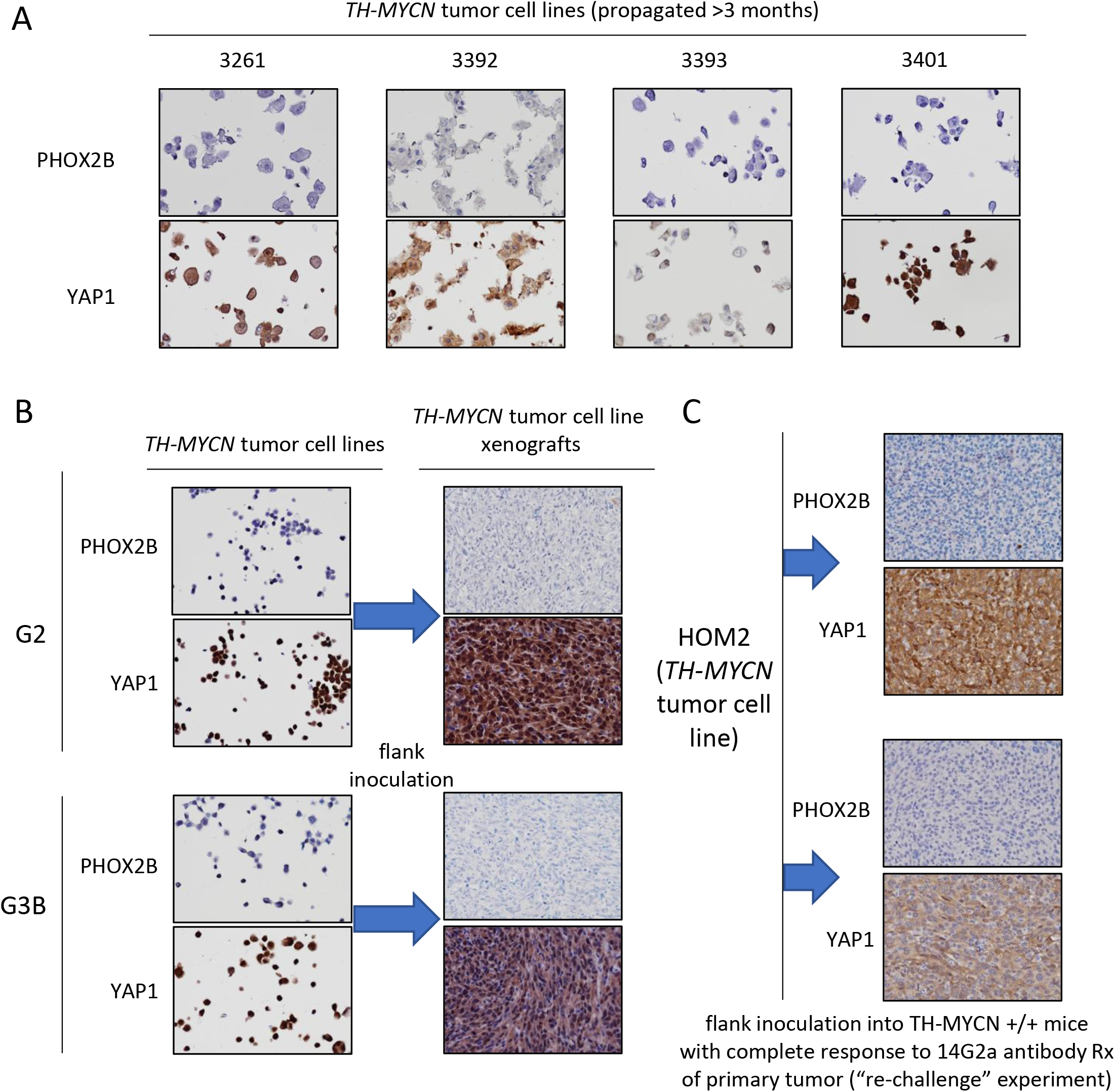
Adrenergic and mesenchymal markers in *TH-MYCN* cell lines and *TH-MYCN* cell line-derived xenografts. (A) Immunocytochemistry for the adrenergic marker, Phox2b, and the mesenchymal marker, Yap1, are shown for a panel of established *TH-MYCN*^+/+^ tumor-derived cell lines, demonstrating near-exclusive mesenchymal expression. (B and C) The same *TH-MYCN*^+/+^ cell line established as a xenograft in the flank of an immunocompromised mouse retains its mesenchymal identity; (B) shows two cell lines as cell line pellets and as xenografted tumors, while (C) shows two xenografts established with a cell line into mice that had a durable complete regression of their autochthonous primary tumors in response to 14G2a therapy (part of the tumor re-challenge trial).

## REFERENCES

1 Coughlan D, Gianferante M, Lynch CF, et al. Treatment and survival of childhood neuroblastoma: Evidence from a population-based study in the United States. Pediatr Hematol Oncol 2017;34:320–30. doi:10.1080/08880018.2017.1373315

2 Hwang WL, Wolfson RL, Niemierko A, et al. Clinical Impact of Tumor Mutational Burden in Neuroblastoma. J Natl Cancer Inst 2019;111:695–9. doi:10.1093/jnci/djy157

3 Noskova H, Kyr M, Pal K, et al. Assessment of Tumor Mutational Burden in Pediatric Tumors by Real-Life Whole-Exome Sequencing and In Silico Simulation of Targeted Gene Panels: How the Choice of Method Could Affect the Clinical Decision? Cancers 2020;12. doi:10.3390/cancers12010230

4 Castriconi R, Dondero A, Bellora F, et al. Neuroblastoma-Derived TGF-β1 Modulates the Chemokine Receptor Repertoire of Human Resting NK Cells. J Immunol 2013;190:5321–8. doi:10.4049/jimmunol.1202693

5 Brandetti E, Veneziani I, Melaiu O, et al. MYCN is an immunosuppressive oncogene dampening the expression of ligands for NK-cell-activating receptors in human high-risk neuroblastoma. OncoImmunology 2017;6:e1316439. doi:10.1080/2162402X.2017.1316439

6 Hashimoto O, Yoshida M, Koma Y, et al. Collaboration of cancer-associated fibroblasts and tumour-associated macrophages for neuroblastoma development: The role of non-tumour stromal cells in neuroblastoma progression. J Pathol 2016;240:211–23. doi:10.1002/path.4769

7 Yu AL, Gilman AL, Ozkaynak MF, et al. Anti-GD2 Antibody with GM-CSF, Interleukin-2, and Isotretinoin for Neuroblastoma. N Engl J Med 2010;363:1324–34. doi:10.1056/NEJMoa0911123

8 Mody R, Naranjo A, Van Ryn C, et al. Irinotecan-temozolomide with temsirolimus or dinutuximab in children with refractory or relapsed neuroblastoma (COG ANBL1221): an open-label, randomised, phase 2 trial. Lancet Oncol 2017;18:946–57. doi:10.1016/S1470-2045(17)30355-8

9 Mody R, Yu AL, Naranjo A, et al. Irinotecan, Temozolomide, and Dinutuximab With GM-CSF in Children With Refractory or Relapsed Neuroblastoma: A Report From the Children’s Oncology Group. J Clin Oncol Off J Am Soc Clin Oncol 2020;38:2160–9. doi:10.1200/JCO.20.00203

10 Hanahan D. Transgenic mice as probes into complex systems. Science 1989;246:1265–75. doi:10.1126/science.2686032

11 Roussel MF, Stripay JL. Modeling pediatric medulloblastoma. Brain Pathol 2019;:bpa.12803. doi:10.1111/bpa.12803

12 Feuring-Buske M, Gerhard B, Cashman J, et al. Improved engraftment of human acute myeloid leukemia progenitor cells in beta 2-microglobulin-deficient NOD/SCID mice and in NOD/SCID mice transgenic for human growth factors. Leukemia 2003;17:760–3. doi:10.1038/sj.leu.2402882

13 Weiss WA. Targeted expression of MYCN causes neuroblastoma in transgenic mice. EMBO J 1997;16:2985–95. doi:10.1093/emboj/16.11.2985

14 Rasmuson A, Segerström L, Nethander M, et al. Tumor Development, Growth Characteristics and Spectrum of Genetic Aberrations in the TH-MYCN Mouse Model of Neuroblastoma. PLoS ONE 2012;7:e51297. doi:10.1371/journal.pone.0051297

15 Hackett CS, Hodgson JG, Law ME, et al. Genome-wide array CGH analysis of murine neuroblastoma reveals distinct genomic aberrations which parallel those in human tumors. Cancer Res 2003;63:5266–73.

16 Teitz T, Stanke JJ, Federico S, et al. Preclinical models for neuroblastoma: establishing a baseline for treatment. PloS One 2011;6:e19133. doi:10.1371/journal.pone.0019133

17 Webb ER, Lanati S, Wareham C, et al. Immune characterization of pre-clinical murine models of neuroblastoma. Sci Rep 2020;10:16695. doi:10.1038/s41598-020-73695-9

18 Kroesen M, Brok IC, Reijnen D, et al. Intra-adrenal murine TH-MYCN neuroblastoma tumors grow more aggressive and exhibit a distinct tumor microenvironment relative to their subcutaneous equivalents. Cancer Immunol Immunother CII 2015;64:563–72. doi: 10.1007/s00262-015-1663-y

19 Kroesen M, Büll C, Gielen PR, et al. Anti-GD2 mAb and Vorinostat synergize in the treatment of neuroblastoma. Oncoimmunology 2016;5:e1164919. doi:10.1080/2162402X.2016.1164919

20 Voeller J, Erbe AK, Slowinski J, et al. Combined innate and adaptive immunotherapy overcomes resistance of immunologically cold syngeneic murine neuroblastoma to checkpoint inhibition. J Immunother Cancer 2019;7:344. doi:10.1186/s40425-019-0823-6

21 Boeva V, Louis-Brennetot C, Peltier A, et al. Heterogeneity of neuroblastoma cell identity defined by transcriptional circuitries. Nat Genet 2017;49:1408–13. doi:10.1038/ng.3921

22 van Groningen T, Koster J, Valentijn LJ, et al. Neuroblastoma is composed of two super-enhancer-associated differentiation states. Nat Genet 2017;49:1261–6. doi:10.1038/ng.3899

23 Haraguchi S, Nakagawara A. A simple PCR method for rapid genotype analysis of the TH-MYCN transgenic mouse. PloS One 2009;4:e6902. doi:10.1371/journal.pone.0006902

24 Hansford LM, Thomas WD, Keating JM, et al. Mechanisms of embryonal tumor initiation: Distinct roles for MycN expression and MYCN amplification. Proc Natl Acad Sci 2004;101:12664–9. doi:10.1073/pnas.0401083101

25 Carlson L-M, Rasmuson A, Idborg H, et al. Low-dose aspirin delays an inflammatory tumor progression in vivo in a transgenic mouse model of neuroblastoma. Carcinogenesis 2013;34:1081–8. doi:10.1093/carcin/bgt009

26 Layer JP, Kronmüller MT, Quast T, et al. Amplification of N-Myc is associated with a T-cell-poor microenvironment in metastatic neuroblastoma restraining interferon pathway activity and chemokine expression. Oncoimmunology 2017;6:e1320626. doi: 10.1080/2162402X.2017.1320626

27 Asgharzadeh S, Salo JA, Ji L, et al. Clinical significance of tumor-associated inflammatory cells in metastatic neuroblastoma. J Clin Oncol Off J Am Soc Clin Oncol 2012;30:3525–32. doi:10.1200/JCO.2011.40.9169

28 Tarek N, Le Luduec J-B, Gallagher MM, et al. Unlicensed NK cells target neuroblastoma following anti-GD2 antibody treatment. J Clin Invest 2012;122:3260–70. doi:10.1172/JCI62749

29 Zingoni A, Molfetta R, Fionda C, et al. NKG2D and Its Ligands: ‘One for All, All for One’. Front Immunol 2018;9:476. doi:10.3389/fimmu.2018.00476

30 Zamora AE, Crawford JC, Thomas PG. Hitting the Target: How T Cells Detect and Eliminate Tumors. J Immunol Baltim Md 1950 2018;200:392–9. doi:10.4049/jimmunol.1701413

31 Theruvath J, Smith B, Linde MH, et al. Abstract PR07: GD2 is a macrophage checkpoint molecule and combined GD2/CD47 blockade results in synergistic effects and tumor clearance in xenograft models of neuroblastoma and osteosarcoma. In: Oral Presentations - Proffered Abstracts. American Association for Cancer Research 2020. PR07–PR07. doi:10.1158/1538-7445.PEDCA19-PR07

32 van Groningen T, Akogul N, Westerhout EM, et al. A NOTCH feed-forward loop drives reprogramming from adrenergic to mesenchymal state in neuroblastoma. Nat Commun 2019;10:1530. doi:10.1038/s41467-019-09470-w

33 Shi H, Tao T, Abraham BJ, et al. ARID1A loss in neuroblastoma promotes the adrenergic-to-mesenchymal transition by regulating enhancer-mediated gene expression. Sci Adv 2020;6:eaaz3440. doi:10.1126/sciadv.aaz3440

34 Huang Y, Tsubota S, Nishio N, et al. Combination of tumor necrosis factor-α and epidermal growth factor induces the adrenergic-to-mesenchymal transdifferentiation in SH-SY5Y neuroblastoma cells. Cancer Sci 2021;112:715–24. doi:10.1111/cas.14760

35 Heiss P, Bermatz S, Wehnes H, et al. Distribution of disialoganglioside GD2-antigen and binding of anti-GD2 antibodies on spheroids of neuroblastoma cell line. Anticancer Res 1997;17:3145–7.

36 Kildisiute G, Kholosy WM, Young MD, et al. Tumor to normal single-cell mRNA comparisons reveal a pan-neuroblastoma cancer cell. Sci Adv 2021;7. doi:10.1126/sciadv.abd3311

37 Wolpaw AJ, Grossmann LD, Dong MM, et al. Epigenetic state determines inflammatory sensing in neuroblastoma. Cancer Biology 2021. doi:10.1101/2021.01.27.428523

38 Wolf BJ, Choi JE, Exley MA. Novel Approaches to Exploiting Invariant NKT Cells in Cancer Immunotherapy. Front Immunol 2018;9:384. doi:10.3389/fimmu.2018.00384

39 Song L, Asgharzadeh S, Salo J, et al. Valpha24-invariant NKT cells mediate antitumor activity via killing of tumor-associated macrophages. J Clin Invest 2009; 119:1524–36. doi:10.1172/JCI37869

40 Mantovani A, Allavena P, Sica A, et al. Cancer-related inflammation. Nature 2008;454:436–44. doi:10.1038/nature07205

41 Sica A, Larghi P, Mancino A, et al. Macrophage polarization in tumour progression. Semin Cancer Biol 2008;18:349–55. doi:10.1016/j.semcancer.2008.03.004

42 Dysthe M, Parihar R. Myeloid-Derived Suppressor Cells in the Tumor Microenvironment. In: Birbrair A, ed. Tumor Microenvironment. Cham::Springer International Publishing 2020. 117–40. doi:10.1007/978-3-030-35723-8_8

43 Trotta R, Dal Col J, Yu J, et al. TGF-beta utilizes SMAD3 to inhibit CD16-mediated IFN-gamma production and antibody-dependent cellular cytotoxicity in human NK cells. J Immunol Baltim Md 1950 2008; 181:3784–92. doi:10.4049/jimmunol.181.6.3784

44 Tran HC, Wan Z, Sheard MA, et al. TGFβR1 Blockade with Galunisertib (LY2157299) Enhances Anti-Neuroblastoma Activity of the Anti-GD2 Antibody Dinutuximab (ch14.18) with Natural Killer Cells. Clin Cancer Res Off J Am Assoc Cancer Res 2017;23:804–13. doi:10.1158/1078-0432.CCR-16-1743

45 Cornel AM, Mimpen IL, Nierkens S. MHC Class I Downregulation in Cancer: Underlying Mechanisms and Potential Targets for Cancer Immunotherapy. Cancers 2020;12:1760. doi:10.3390/cancers12071760

46 Norris MD, Burkhart CA, Marshall GM, et al. Expression of N-myc and MRP genes and their relationship to N-myc gene dosage and tumor formation in a murine neuroblastoma model. Med Pediatr Oncol 2000;35:585–9. doi:10.1002/1096-911x(20001201)35:6<585::aid-mpo20>3.0.co;2-p

47 Moore HC, Wood KM, Jackson MS, et al. Histological profile of tumours from MYCN transgenic mice. J Clin Pathol 2008;61:1098–103. doi:10.1136/jcp.2007.054627

48 Nguyen R, Moustaki A, Norrie JL, et al. Interleukin-15 Enhances Anti-GD2 Antibody-Mediated Cytotoxicity in an Orthotopic PDX Model of Neuroblastoma. Clin Cancer Res Off J Am Assoc Cancer Res 2019;25:7554–64. doi:10.1158/1078-0432.CCR-19-1045

49 Barry WE, Jackson JR, Asuelime GE, et al. Activated Natural Killer Cells in Combination with Anti-GD2 Antibody Dinutuximab Improve Survival of Mice after Surgical Resection of Primary Neuroblastoma. Clin Cancer Res Off J Am Assoc Cancer Res 2019;25:325–33. doi:10.1158/1078-0432.CCR-18-1317

