## Supplementary material for "Anti-GD2 antibody therapy alters the neuroblastoma tumor microenvironment and extends survival in *TH-MYCN* mice": GD2SuppMethodsFile

**SUPPLEMENTAL METHODS:**

**Human neuroblastoma cell lines**. Patient-derived neuroblastoma cell lines COGN603, CHLA122, IMR5, NLF, CHLA20, SKNBE1, SKNBE2C, CHLA95, CHLA108, SHEP, and COGN203 were obtained from the cell line bank at the Children’s Hospital of Philadelphia. Many were previously received from C. Patrick Reynolds (Texas Tech University of Health Sciences; cccells.org). Cells were thawed, plated in complete RPMI (10% FBS + 1% Penicillin-Streptomycin + 1% L-glutamine), and passaged at ~80% confluency using Versene solution, 0.48mM EDTA (Gibco). IMR5 (adrenergic state) and SHEP (mesenchymal state) cells were used as controls in assessing lineage-defining proteins.

**Primary anti-mouse and anti-human antibodies used in flow cytometry**

Primary anti-mouse and anti-human antibodies specific for the following antigens were purchased from BioLegend (alternate names in parentheses; clone names in brackets): CD3 [17A2], CD45 [30-F11], NK1.1 [PK136], CD335 (NKp46) [29A1.4], Ly49A [YE1/48.10.6], CD314 (NKG2D) [CX5], CD159a (NKG2A) [16A11], GD2 [14G2a], MHC I (H-2K/H-2D) [28-8-6], Rae-1γ [CX1], TGF-1 (LAP) [TW4-2F8], CD11b [M1/70], CD11c [N418], F4/80 [BM8], Ly-6G/Ly-6C (Gr-1) [RB6-8C5], Ly-6G [1A8], CD38 [90], CD107a (LAMP-1) [1D4B], TCR γδ [GL3], CD56 [HCD56]. Additional antibodies include Ly49H [3D10], and MHC Class II (I-A/I-E) [M5/114.15.2] from eBioscience, IL-15Rα [888220], ULBP-1 [237104] and NCAM1 [809220] from R&D Systems, TGFβR-1 [RM0016-3A11] from Signalway, and ULBP-1 from ThermoFisher Scientific. PBS57-loaded CD1d tetramers were obtained from the NIH Tetramer Facility (Atlanta, GA).

**Flow analysis and gating strategy**. FCS files were analyzed using FlowJo software (Tree Star: Ashland OR). *TH-MYCN* neuroblastoma cell lines were gated by selecting live single CD45^-^/NCAM1^+^ cells while human neuroblastoma cell lines were gated by selecting live single CD45^-^/CD56^+^ cells. Tumors from mice treated with 14G2a, isotype-control antibody, and PBS were analyzed using the same gating strategy. Frequencies of lymphoid and myeloid cells were determined from live doublet-excluded CD45^+^ parent cell populations. Among the lymphocyte populations, NK cell frequencies were defined as CD3^-^/NK1.1^+^/NKp46^+^ cells, T-cell frequencies were defined as CD3^+^/NK1.1^-^/NKp46^-^ cells, invariant natural killer T cells were defined as CD1d tetramer-binding/CD3^+^, and γδ T cells were defined as γδ TCR^+^/CD3^+^ cells. NK cell surface expression of activating (Ly49H, IL15Rα, and NKG2D) and inhibitory (Ly49A, NKG2A, and TGFβR-1) receptors were assessed. Among the myeloid populations, dendritic cells (DCs) were defined as F4/80^-^/CD11c^+^/CD11b^+^, monocytic myeloid-derived suppressor cells (M-MDSCs) were defined as Gr-1^+^/CD11b^+^/Ly6G^-^, granulocytic myeloid-derived suppressor cells (G-MDSCs) were defined as Gr-1^+^/CD11b^+^/Ly6G^+^, and macrophages were defined as MHC-II^+^/F4/80^+^/CD11c^-^.

**Detection of lineage-defining proteins.** For immunoblots, 30ug of protein lysate were electrophoresed through a 4-12% Bis-Tris NuPAGE Gel and transferred to PVDF membranes using an iBlot Gel Transfer device. Membranes were blocked for 60 min with a 5% milk in 0.1% Tween-20 in TBS (TTBS) solution followed by TTBS wash, then incubated overnight at 4°C with antibodies for Phox2B (1:1000; Santa Cruz Biotechnology, clone B-11), Yap1 (1:1000; Cell Signaling, polyclonal) or β-tubulin (Sigma Aldrich, clone AA2). Secondary antibody (1:1000-1:2000) was applied for 60-120 min at room temperature, followed by wash steps and detection via horseradish peroxidase chemiluminescent substrate (Millipore). For testing cell lines by immunocytology, 10-20 x 10^6^ live cells were suspended in a 1% agarose solution in isosmotic PBS at 50C° and transferred to a 1.5 Eppendorf tube and placed on ice until solid, then removed and fixed with formalin. For testing primary *TH-MYCN* tumors or tumor cell line xenografts, tumors were removed in terminal surgeries and formalin-fixed and paraffin embedded. Slides of FFPE materials were deparaffinized, blocked with 5% horse serum, and stained. Yap1 (Cell Signaling Technology, clone D8H1X) and Phox2B (Abcam, clone EPR14423) were both used at 1:50. Slides were counterstained with hematoxylin. Images were captured using a Nikon 80i Upright microscope using the 10x and 20x lenses.
